# Investigation of latent representation of toxicopathological images extracted by CNN model for understanding compound properties *in vivo*

**DOI:** 10.1101/2023.07.23.550191

**Authors:** Shotaro Maedera, Tadahaya Mizuno, Hiroyuki Kusuhara

## Abstract

Toxicopathological images acquired during safety assessment elucidate an individual’s biological responses to a given compound, and their numerization can yield valuable insights contributing to the assessment of compound properties. Currently, toxicopathological images are mainly encoded as pathological findings, evaluated by pathologists, which introduces challenges when used as input for modeling, specifically in terms of representation capability and comparability. In this study, we assessed the usefulness of latent representations extracted from toxicopathological images using Convolutional Neural Network (CNN) in estimating compound properties *in vivo*. Special emphasis was placed on examining the impact of learning pathological findings, the depth of frozen layers during learning, and the selection of the layer for latent representation. Our findings demonstrate that a machine learning model fed with the latent representation as input surpassed the performance of a model directly employing pathological findings as input, particularly in the classification of a compound’s Mechanism of Action and in predicting late-phase findings from early-phase images in repeated-dose tests. While learning pathological findings did improve accuracy, the magnitude of improvement was relatively modest. Similarly, the effect of freezing layers during learning was also limited. Notably, the selection of the layer for latent representation had a substantial impact on the accurate estimation of compound properties *in vivo*.

## 1 Introduction

Given the frequency of drug withdrawal due to unforeseen adverse reactions and the potential for drug repositioning based on novel pharmacological effects, it is inferred that low molecular weight compounds, such as drugs, possess multiple and unrecognized effects [1], [2]. These effects can be described as the biological responses of an organism to the compound [3], [4]. Numerization and subsequent data-driven analysis are valuable in understanding compound effects, including toxicity prediction [5], [6]. Pathological imaging, specifically toxicopathology, plays a crucial role in safety assessment of compounds, providing a repository of in vivo biological responses from experimental animals exposed to specific compounds [7]. However, toxicopathological images are predominantly evaluated based on manually crafted pathological findings, which may have limited dimensionality and subjectivity, although these pathological findings offer notable advantages in terms of interpretability and acceptance due to accumulated knowledge, thereby potentially hindering their utility in subsequent data analysis [8], [9].

Representation learning, a specialized paradigm using neural networks for feature extraction [10], allows for meaningful information extraction from variable-length images, converting them into fixed-length data representations [11], [12]. While representation learning has shown advancements in cancer diagnosis and prognosis prediction, its application in toxicopathological images is limited [13]. Thus, this study assessed the utility of latent representations derived from a Convolutional Neural Network (CNN) for estimating compound properties *in vivo*, focusing on toxicopathological images of the liver due to its first-pass effect during oral administration [14].

In this study, we employed the intermediate layer of a Convolutional Neural Network (CNN) that was initially pre-trained on ImageNet as the latent representation. This network was fine-tuned for an auxiliary task specific to the domain at hand, which is a common approach in image representation learning [15], [16]. Our focus was on pathological finding prediction as the auxiliary task, leading us to investigate three key aspects: (1) whether the CNN models could learn pathological findings of toxicopathological images used in safety assessment, (2) which layer should be frozen during the learning of pathological findings, and (3) which layer should be used for feature extraction as latent representation. To conduct this investigation, we fine-tuned the pre-trained CNN models on ImageNet using a substantial dataset of toxicopathological images containing pathological findings from Open TG-GATEs (TGGATEs). The fine-tuning process encompassed all possible combinations of frozen layers and representation layers (**Figure 1a**) [17]. Subsequently, we extracted latent representations of toxicopathological images treated with various compounds using each constructed model. These representations were then evaluated based on the performance of downstream tasks related to compound effects *in vivo*, including classifying the mechanism of action (MoA) and predicting late-phase findings from early-phase images in repeated-dose tests.

**Figure 1.**
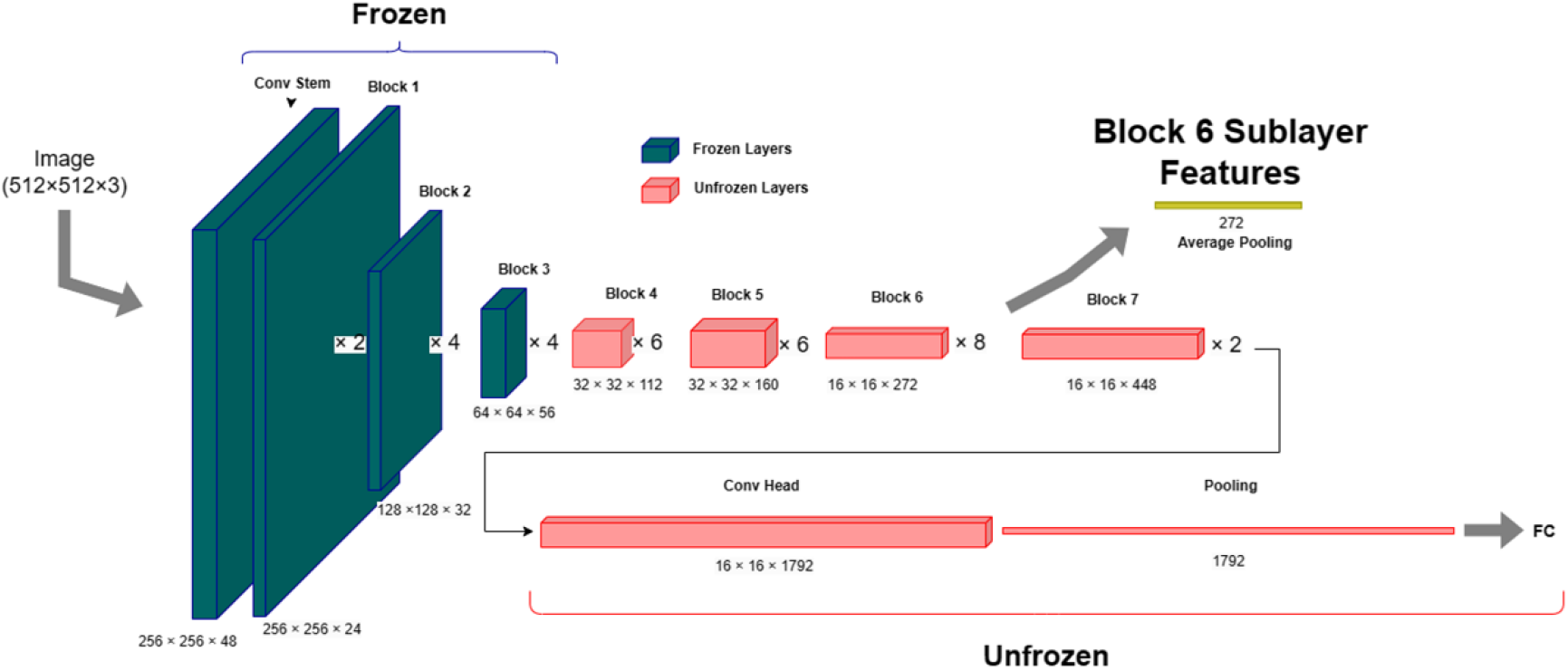
Illustration of CNN model mainly employed in this study. We prepared this illustration using draw.io (https://app.diagrams.net/).

## 2 Related Work

Concerning image feature extraction, the Scale-invariant feature transform (SIFT) has been a widely utilized method since the early 2000s [18]. SIFT features have demonstrated remarkable robustness in matching across a significant range of affine distortion, changes in 3D viewpoint, addition of noise, and alterations in illumination conditions [19].

Since the introduction of AlexNet in 2012, feature extraction methods employing the intermediate layer of CNNs have been proposed and remain a popular approach for image representation learning [19]. Through the training of CNNs on annotated image datasets like ImageNet or extensive datasets such as JFT-300M, it has become feasible to achieve high-quality feature extraction for numerous computer vision tasks. Notable examples of CNN architectures include ResNet, the pioneering CNN that addresses the vanishing gradient problem through skip connections, and EfficientNet, an outcome of research focused on efficient model scaling [20], [21]. Other prominent neural network-based techniques utilize the transformer architecture, which exclusively employs attention mechanisms instead of recurrent neural networks (RNNs) and CNNs [22]. Initially, the transformer architecture was primarily employed as a natural language processing model. However, in 2020, the Vision Transformer (ViT) was introduced, which utilizes transformers for image processing by tokenizing image segments [23]. These transformer-based image recognition models can also serve as feature extractors. In recent years, researchers have been actively working on developing self-supervised learning methods as well [24]–[27].

Representation learning has previously been applied to pathological images, and its potential application to cancer diagnosis is currently being explored [28], [29]. In the simple yet representative approach that involves utilizing the intermediate layer of a CNN pretrained on ImageNet, the significance of determining the appropriate layers for feature extraction and the choice of auxiliary tasks to be learned is under verification [30], [31]. For instance, utilizing intermediate layers is generally favored over using the final layer in the majority of cases, as the latter tends to be more task-specific. However, the specific layer selection depends on the particular task at hand and the same holds true for auxiliary tasks. Notably, there is limited research on representation learning of toxicopathological images. In 2021, Hoefling et al. and in 2022, Koohbanani et al. published works related to toxicopathology [32], [33]. However, both studies primarily focused on anomaly detection using normal rat or mouse tissues, without a specific emphasis on pathological findings [13]. Therefore, the research questions are twofold: (1) whether latent representation of toxicopathological images is useful for understanding compound effects *in vivo* as input for data analysis and (2) which factor affects representation learning models for extracting good representations for that purpose.

## 3 Methods

### 3.1 Computational resource and environment

The PyTorch version 1.12.1 (https://pytorch.org/) was selected as the deep learning framework in this study. We conducted all deep learning and feature extraction operations using a GeForce RTX 3090 (NVidia).

### 3.2 Data preparation

We collected a total of 23,724 pathological images of the liver from the Open TG-GATEs (TGGATEs) database, along with corresponding pathological finding type annotations and individual rat information from TogoDB (http://togodb.biosciencedbc.jp/togodb/view/open_tggates_main#en). Mechanism of Action (MoA) annotations were obtained from DrugBank (https://go.drugbank.com/).

For efficient identification of the organ area within each Whole Slide Image (WSI), we employed a specific method. Initially, we compressed each WSI to 1/64 of its original size. Subsequently, a pixel was considered blank if its average 24-bit RGB value exceeded 200. The compressed image was then divided into a grid of 16×16 pixel squares. Squares containing over 60% non-blank pixels were identified as organ area squares. From these squares, we randomly selected 100 small images. In the subsequent analysis, a 512×512 pixel area from the center of each square was cropped and used as a patch. However, when employing “random crop” as the data augmentation method for deep learning, a 512×512 pixel area was randomly cropped from each square during the learning process. These operations were performed using OpenCV-Python version 4.6.0.66 (https://pypi.org/project/opencv-python/) and OpenSlide-Python version 1.2.0 (https://openslide.org/). The meta information of TGGATEs is available at http://togodb.biosciencedbc.jp and is summarized in **Supplementary Figure 1**.

### 3.3 Pathological finding selection utilizing weakly supervised learning

We employed weakly supervised learning (WSL) to identify widely distributed and easily recognizable pathological findings in Whole Slide Images (WSI) and annotated the patches accordingly. Out of a total of 65 pathological findings, we excluded the “Dead” finding, which does not pertain to the status of pathological images. For each of the remaining 64 pathological findings, we trained EfficientNetB4 using WSI-level finding type annotations directly for patches. The TGGATEs images were divided into two groups, namely the TGGATEs WSL train group and TGGATEs WSL validation group, with a 4:1 ratio, ensuring that images from the same experimental group belonged to the same group.

Certain pathological findings were not observed in both the WSL train group and validation group with respect to the Experimental Group, leading us to exclude them (refer to **Supplementary Figure 1** for Experimental Group). Subsequently, EfficientNetB4 models were constructed for the remaining 44 pathological findings (**Supplementary Table 1**). The deep learning settings are described in **Supplementary Note 1**. After the training process, we predicted the probability of each patch having the respective pathological finding. Eye observation was used to evaluate the homogeneity of randomly sampled patches, resulting in the exclusion of 28 findings. Based on the homogeneity of patches with the top 30 and worst 30 probabilities, we selected eight pathological findings (**Table 1**, **Supplementary Figure 2**). It is important to note that two different researchers, S.M. and T.M., independently conducted the eye observation. We extracted patches with any of the eight selected finding types with a probability of 0.5 or more. Additionally, an equal number of patches were obtained from WSIs where no finding types were present. These patches were used for the model construction.

**Table 1.** List of pathological findings mainly investigated in this study.

| Finding Types | Train |  |  | Validation |  |  | Test |  |  |
| --- | --- | --- | --- | --- | --- | --- | --- | --- | --- |
|  | posi | nega | ratio | posi | nega | ratio | posi | nega | ratio |
| Proliferation, bile duct | 5067 | 46296 | 9.87% | 903 | 16218 | 5.27% | 4722 | 32946 | 12.54% |
| Ground glass appearance | 4907 | 46456 | 9.55% | 1191 | 15930 | 6.96% | 3841 | 33827 | 10.20% |
| Increased mitosis | 5033 | 46330 | 9.80% | 1564 | 15557 | 9.14% | 3448 | 34220 | 9.15% |
| Inclusion body, intracytoplasmic | 195 | 51168 | 0.38% | 171 | 16950 | 1.00% | 496 | 37172 | 1.32% |
| Deposit, pigment | 1789 | 49574 | 3.48% | 500 | 16621 | 2.92% | 603 | 37065 | 1.60% |
| Single cell necrosis | 2892 | 48471 | 5.63% | 561 | 16560 | 3.28% | 3696 | 33972 | 9.81% |
| Vacuolization, cytoplasmic | 8142 | 43221 | 15.85% | 2537 | 14584 | 14.82% | 3979 | 33689 | 10.56% |
| Swelling | 4207 | 47156 | 8.19% | 1952 | 15169 | 11.40% | 3155 | 34513 | 8.38% |

### 3.4 TGGATEs data split

To ensure careful data structure in TGGATEs, we performed data splitting using a specific procedure (**Supplementary Figure 3**). Initially, the patches selected in the previous process were divided into five folds using the repeated stratified group K-fold method. This division was carried out to satisfy two conditions: (1) Equal representation of each selected pathological finding in each fold and (2) Patches of rats treated with the same compound belonging to the same fold. The term “repeated” indicates that the algorithm attempts multiple times to achieve such a split Next, the dataset resulting from combining four parts of the fold created in the previous step was again split into five folds using the repeated stratified group K-fold method. This time, patches were divided so that rats belonging to the same experimental group were present in the same fold. We combined the following folds: one from the first split, which was not used in the second split, and the other from the second split. These were named the TGGATEs test groups. This procedure enabled the creation of test data that allowed us to evaluate generalization performance, rather than overfitting to features specific to the administered compounds or experimental groups.

For patches not included in the test dataset, one of the folds created during the second data split was designated as the TGGATEs validation group, and the remaining three folds were labeled as the TGGATEs train group. All experiments were performed with 5 different seeds, and the results were reported as the mean and standard deviation.

### 3.5 Extraction of latent representation of toxicopathological images

To extract the latent representation of toxicopathological images, we followed two steps. Firstly, we performed global average pooling on the feature map of the targeted layer to obtain a feature vector at the patch level [34]. Subsequently, we defined the latent representation of a toxicopathological image that describes a compound of interest as a WSI-level feature. This was achieved by taking the arithmetic mean of the patch-level features derived from the sampled patches within the WSI.

In this study, we examined the optimal layer for freezing during training and for feature extraction to investigate the latent representations of toxicopathological images. We froze several layers during training, as depicted in **Figure 1**, and prepared 10 models. For latent representation extraction, we compared all nine layers in the EfficientNetB4 backbone. Consequently, a total of 90 (10 × 9) different representations were generated for each toxicopathological image.

### 3.6 Leave one out experiment for testing generalization performance

EfficientNetB4 was trained on seven of the eight selected pathological findings, with one finding excluded. The training conditions are described in **Supplementary Note 2**. Subsequently, features were extracted from the patches in the test dataset using both the model we constructed and an ImageNet pretrained model as a control (Control model). For the linear evaluation of latent representations, logistic regression was conducted using the extracted features as explanatory variables and the pathological finding not used in model construction as the dependent variable. L2 regularization with λ set to 1.0 (default setting of scikit-learn module) was applied to the logistic regression model using scikit-learn version 1.0.2.

The evaluation of the logistic regression model followed a specific procedure. The test dataset was divided into five folds using the group K-fold method. This division ensured that patches belonging to the same WSI were present in the same fold. The logistic regression model was trained with four folds and predictions were evaluated using the remaining one fold. Area Under the Receiver Operating Characteristic Curve (AUROC) and mean Average Precision (mAP) were used as evaluation metrics. This process was repeated for all selected pathological findings.

After the LOO experiment, models were prepared, learning from all eight selected pathological findings, which were subsequently employed for further analyses. All results are summarized in **Supplementary Data 1**.

### 3.7 Classification of MoA

We focused on seven MoAs out of a total of 1,394 MoAs in DrugBank, considering data imbalance (**Table 2, Supplementary Table 2**). We employed WSIs that met the following four conditions: (1) WSIs of rats that received compounds with the selected MoA (30 compounds in total), (2) WSIs of rats sampled after four days of repeated administration, (3) WSIs of rats that did not belong to the train or validation groups, and (4) WSIs of rats treated with compounds at concentrations higher than 0. The second and fourth conditions were set to ensure sufficient effects of the dosed compounds on the rats. Some data in TGGATEs involved rats that only received vehicles as a control group, i.e., dosing was performed at zero concentration. The latent representations were used to predict the MoA using multiclass logistic regression with L2 regularization. The experimental design is detailed in **Supplementary Figure 4**. Additionally, we constructed a logistic regression model with all 65 pathological findings as explanatory variables. To assess the model’s performance, we conducted 10 trials of cross-validation, dividing the analysis target into five parts while ensuring that the same compound did not cross the fold. The primary evaluation metric was balanced accuracy, considering the multi-class classification task.

**Table 2.** List of Mechanism of Actions mainly investigated in this study.

| Mechanism of Action | Name of Compounds |
| --- | --- |
| Cyclooxygenase inhibitor | naproxen; sulindac; sulfasalazine; diclofenac; acetaminophen; phenylbutazone; mefenamic acid; indomethacin; aspirin |
| DNA inhibitor | nitrofurantoin; carboplatin; azathioprine; nitrofurazone; lomustine; cyclophosphamide |
| Bacterial 70S ribosome inhibitor | erythromycin ethylsuccinate; chloramphenicol; tetracycline |
| Histamine H2 receptor antagonist | cimetidine; famotidine; ranitidine |
| Peroxisome proliferator-activated receptor alpha agonist | fenofibrate; gemfibrozil; clofibrate |
| Serotonin 2a (5-HT2a) receptor antagonist | chlorpromazine; thioridazine; haloperidol |
| Sulfonylurea receptor 1, Kir6.2 blocker | glibenclamide; tolbutamide; chlorpropamide |

### 3.8 Prediction of the late-phase pathological findings from the early-phase images in repeated-dose tests

TGGATEs contains WSIs from repeated-dose tests, as illustrated in **Supplementary Figure 5**. Using this data, we evaluated the performance of latent representations for predicting late-phase pathological findings from early-phase images (Early-to-late prediction). To conduct the evaluation, we selected WSIs that met the following three conditions: (1) WSIs of rats sampled after four days of repeated administration, (2) WSIs of rats that did not belong to the train or validation groups, and (3) WSIs of rats treated with compounds at concentrations higher than zero. We divided the rats sampled four and eight days after the start of daily administration into the early group, and the rats sampled 15 and 29 days after the start of compound administration into the late group.

During the training phase, we constructed a logistic regression model with L2 regularization, using the latent representations of toxicopathological images from the late group as explanatory variables and the corresponding pathological findings as the objective variable. We selected 39 pathological findings out of a total of 65 findings, based on the availability of a sufficient number of positive images for analysis. In other words, we ensured that positive images existed for both the early and late groups, respectively.

In the inference phase, we calculated predicted values using the logistic regression model constructed above, taking the latent representations of early group images as input. Additionally, we directly used the annotations of each pathological finding as predictions (logistic regression model as a constant function). Each WSI of rat liver treated with a compound in the early group was labeled based on whether a pathological finding was found in at least one rat in the late group that received the same compound as the rat to which the WSI belonged. WSIs were labeled positive if the finding was present and negative if it was absent. Using these labels, each prediction was evaluated with AUROC and mAP, which do not require a threshold.

## 4 Results and Discussion

### 4.1 Construction of representation learning models of toxicopathological images by learning pathological findings

In this study, we utilized the toxicological dataset provided by TGGATEs, which contains Hematoxylin and Eosin stained images of rat liver. The dataset comprises daily administration of 160 chemicals at multiple concentrations, and liver pathological images were obtained at various time points. To simplify the training process, we focused on selecting pathological findings that are easily manageable in patch-based analysis, commonly employed in pathological image analysis. We prepared weakly supervised learners by providing WSI labels to the constituent patches and selected pathological findings that weak learners could easily judge at the patch-level (**Supplementary Figure 2**).

To assess the hypothesis that learning pathological findings used in safety assessment improves the generalization performance of models in extracting information from toxicopathological images, we conducted leave-one-out (LOO) experiments for each pathological finding. Specifically, we chose one pathological finding, fine-tuned a CNN model pretrained on ImageNet with classification tasks of pathological findings other than the selected one, and obtained a representation learning model. Note that EfficientNetB4 was primarily used in this study, and some results were confirmed using ResNet50 as well (**Supplementary Note 3**). We constructed a logistic regression model for the selected finding using the latent representation, a fixed-length vector obtained with the model, and evaluated its performance.

The results were summarized by mean of findings and visualized as a heatmap in **Figure 2** and **Supplementary Figure 4**. Most of the fine-tuned models exhibited slightly higher scores compared to the corresponding pretrained (control row in Figures) models in each feature (columns) (**Figure 2a and b**). Regarding the layer dependence, in the column-wise analysis, we discussed which layer should be frozen during fine-tuning on pathological findings, i.e., how deep pathological findings affect the model. In the row-wise analysis, we explored which layer of feature map should be employed for the latent representation of compound effects *in vivo*. The median of column-wise variances was smaller than that of the row-wise, suggesting that the selection of the layer for feature extraction has a greater impact on performance than the depth of learning pathological findings (**Figure 2c and d**). Note that although the score heatmap patterns were not entirely consistent across findings, the overall tendencies were similar (**Supplementary Figure 6**).

**Figure 2.**
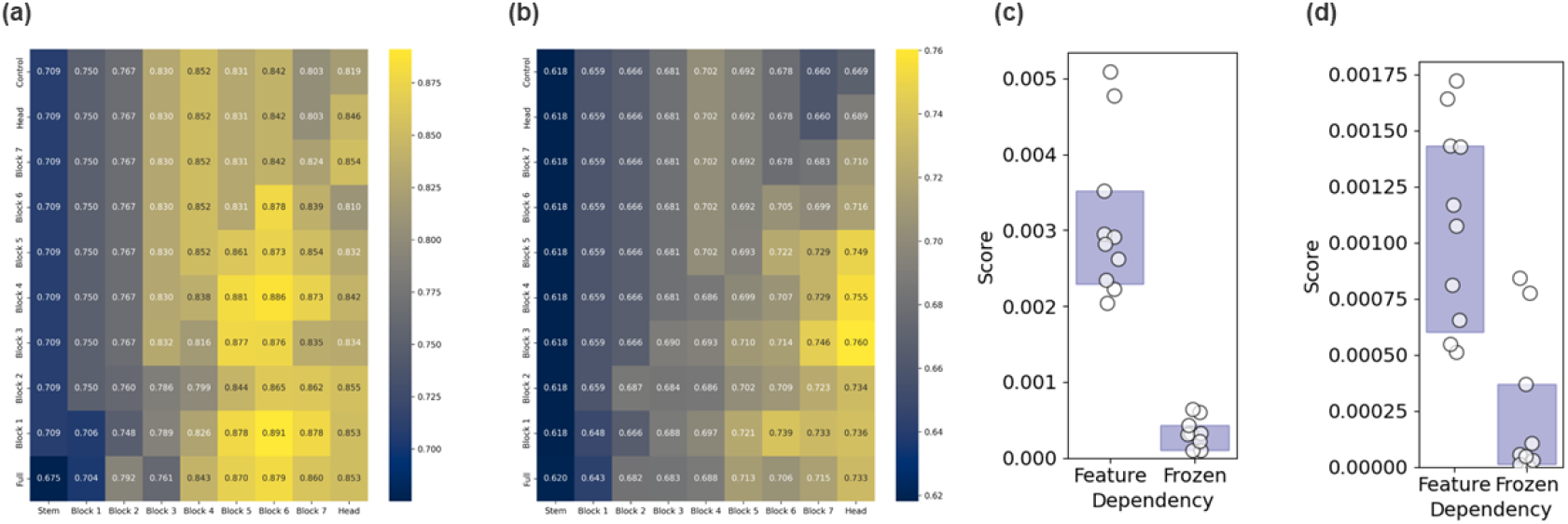
Construction of representation learning models of toxicopathological images by learning pathological findings. (a, b) Heatmaps depicting the averaged AUROC and mAP values. Each heatmap illustrates the performance of finding prediction, with the mean values from 5 different seeds visualized. (c, d) Exploration of layer dependency by comparing AUROC and mAP variances across different layers. The heatmap presents the median values of these metrics for the frozen layer (column-wise) and the feature layer (row-wise) and is represented by open symbols. The filled box denotes the upper and lower 95% confidence intervals, calculated using the bias-corrected and accelerated bootstrap method.

These results indicate that learning pathological findings utilized in safety assessment improves the generalization performance of latent representations for pathological finding detection, but to a limited extent. The key to performance lies in selecting the appropriate layer for the representations. For further analyses, we prepared representation learning models trained on all the selected pathological findings simultaneously, with various combinations (90 in total) of frozen layers and feature layers. These models were then used in subsequent analyses as numerization models to understand compound effects *in vivo*.

### 4.2 Classification of MoA using latent representation of toxicopathological images

Next, we attempted to estimate the MoA of compounds administered to rats using the latent representations extracted by the models described earlier. We specifically focused on seven MoAs that were shared by three or more TGGATEs compounds (**Supplementary Table 2**). To simplify the process, we collected pathological images from rats sacrificed on or after the 4th day after repeated dosing of compounds associated with the targeted MoAs, while excluding rats belonging to the train or validation groups (see section 3.7 for details). We utilized the latent representations as input and estimated the MoA using logistic regression, evaluating the performance through five-fold cross-validation. The dataset was divided so that WSIs from the same experimental group belonged to only one fold (**Supplementary Figure 3**).

The results were summarized by the mean of MoAs and visualized as a heatmap in **Figure 3** and **Supplementary Figure 7**. Concerning the effects of learning pathological findings, most of the fine-tuned models exhibited slightly higher scores than the corresponding pretrained (control row in Figures) models in each feature (columns), although the degree of improvement was modest (**Figure 3a**). Regarding the layer dependence, the median of column-wise variances was smaller than that of the row-wise, indicating that the impact of selecting the layer for feature extraction is more significant than how deep learning pathological findings affect the model, consistent with the results of the LOO experiments for pathological finding prediction (**Figure 3b**).

**Figure 3.**
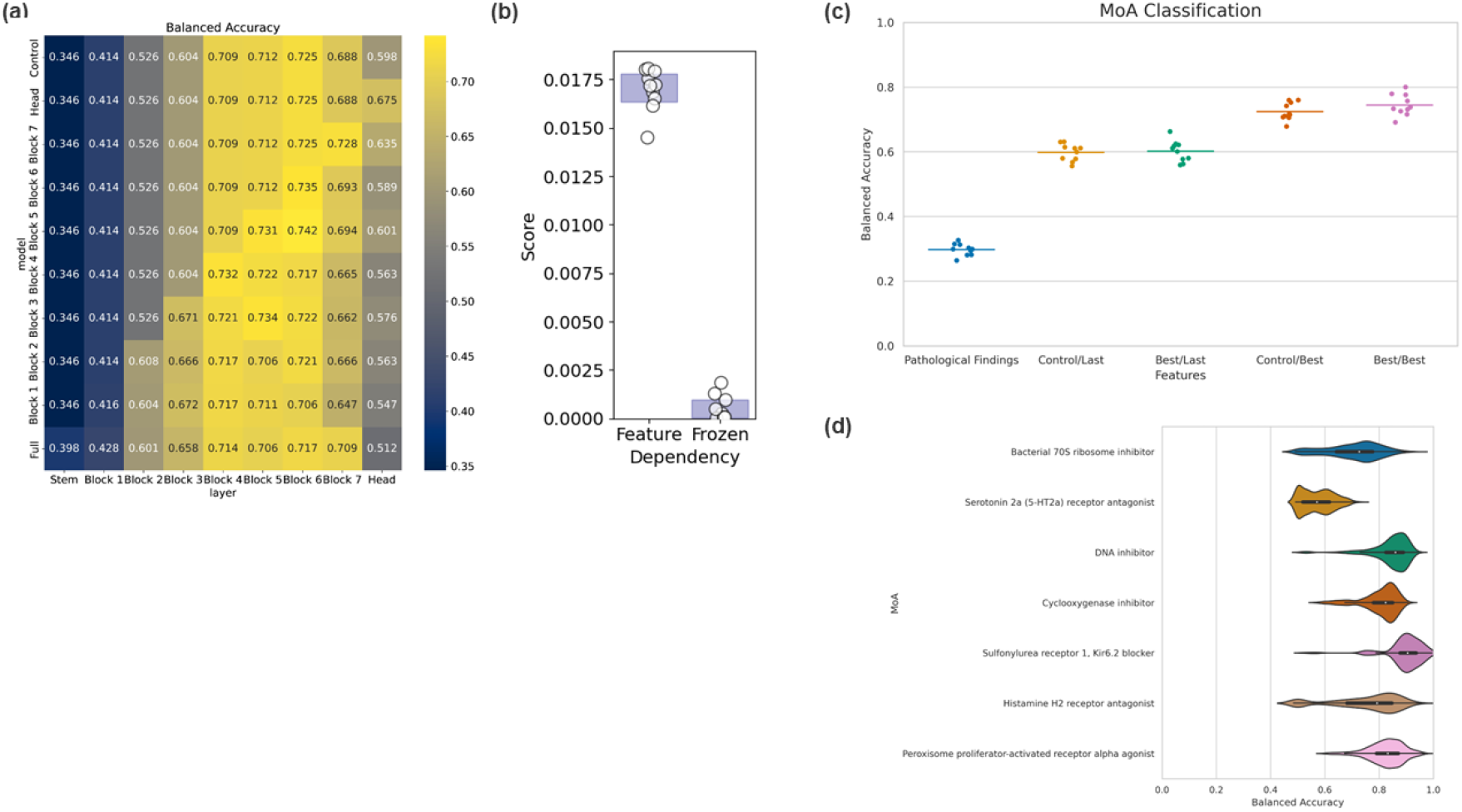
Classification of MoA using latent representation of toxicopathological images. (a) Heatmap displaying the mean Balanced Accuracy of each MoA classification, calculated from 5 different seeds. (b) Strip-plot investigating layer dependency by comparing the variances of Balanced Accuracy for each layer. The medians of the metric in column-wise and row-wise represent the frozen layer and feature layer dependencies, respectively, visualized as open symbols. The filled box shows the upper and lower 95% confidence intervals calculated by the bias-corrected and accelerated bootstrap method. (c) Strip-plot comparing classification performances of representative models. The best combination of the frozen and feature layers, along with the corresponding control, and classification directly using pathological findings as input were selected. The mean Balanced Accuracy of selected MoAs is calculated for 10 cross-validation trials and visualized as points and bars, indicating each fold’s value and mean value, respectively. X/Y represents the Y layer of the X model used for MoA classification. (d) Violin plot presenting the Balanced Accuracy of each MoA.

For interpretability, we selected the best combination of the frozen and feature layer, along with the corresponding control, and compared them to the results of classification directly using pathological findings as input. Notably, the latent representations exhibited superior MoA prediction ability compared to the binary vectors composed of pathological findings, regardless of training on pathological findings (**Figure 3c, Supplementary Figure 8**). This indicates that even ImageNet pre-trained models can capture features related to MoA, which aligns with previous reports in the field of cancer pathology [29]. Additionally, it is worth noting that MoAs such as “5-HT2a antagonist,” which may have minimal direct effects on the liver based on its drug target, were more challenging to predict, while MoAs that could directly affect the liver, such as “DNA inhibitor” due to its broad spectrum, were relatively easier to predict (**Figure 3d**).

### 4.3 Predicting pathological findings in the late-phase of repeated-dose tests using the latent representation of the early-phase images

We examined the predictive capability of latent representations from constructed models for safety assessment. Rats in test groups were divided into early and late groups based on the time of sacrifice. The task involved predicting late-phase pathological findings using early-phase latent representations, referred to as Early-to-Late prediction (**Figure 4a**). Logistic regression models were constructed with late-phase latent representations as explanatory variables, targeting 39 pathological findings observed in sufficient WSIs in TGGATEs (as in section 3.8). Probabilities of pathological findings for early-phase images were obtained using these models. Consistency between predicted probabilities and late-phase pathological findings was assessed with multiple metrics. Results were summarized using a heatmap in **Figure 4** and **Supplementary Figure 9**.

**Figure 4.**
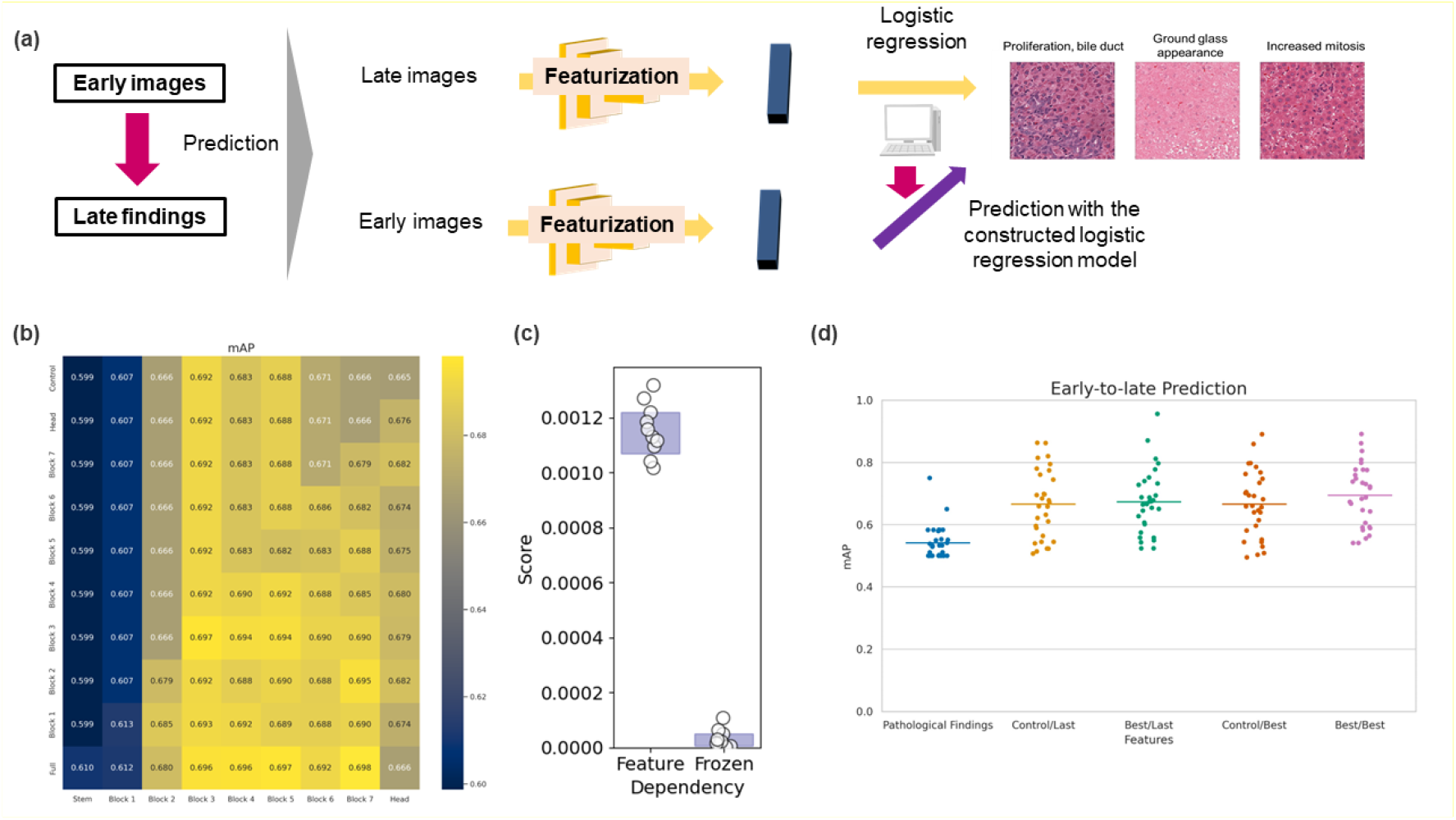
Predicting late-phase pathological findings through latent representation of early-phase images in repeated-dose tests. (a) Illustration of early-to-late prediction of pathological findings. (b) Heatmap displaying the mean mAP of each finding prediction, calculated from 5 different seeds. (c) Strip-plot investigating layer dependency by comparing the variances of mAP for each layer. The medians of mAP in column-wise and row-wise represent the frozen layer and feature layer dependencies, respectively, visualized as open symbols. The filled box shows the upper and lower 95% confidence intervals calculated using the bias-corrected and accelerated bootstrap method. (d) Strip-plot comparing prediction performances of representative models. The best combination of the frozen and feature layers, along with the corresponding control, and classification directly using pathological findings as input were selected. The mAP of the models is visualized as points and bars, indicating each finding’s value and mean value, respectively. X/Y represents the Y layer of the X model used for early-to-late prediction.

Regarding learning pathological findings, most fine-tuned models exhibited slightly higher scores than their corresponding pretrained models (control row in Figures) for each feature (columns) (**Figure 4b**). Layer dependence analysis revealed that the impact of selecting the layer for feature extraction was greater than the depth of pathological findings affecting the model, consistent with other task results (**Figure 4c**). For interpretability, the best combination of frozen and feature layers, along with corresponding controls, was identified. Classification using pathological findings directly as input was also performed. Remarkably, latent representations demonstrated superior predictive performance compared to findings judged by pathologists, regardless of the training pathological findings (**Figure 4d**). These results indicate that representation learning models can anticipate the expression of pathological findings earlier than human observation. It is noteworthy that this ability of CNN models’ latent representations is not limited to those learning pathological findings when an appropriate layer is used for feature extraction.

### 4.4 Evaluation of the relationship between the accuracy of estimating pathological findings and downstream tasks

Lastly, we explored the association between the accuracy of estimating pathological findings during the construction of representation learning models and downstream tasks utilizing the latent representations. The correlation of compound property prediction performances among the tasks was computed using the prepared models. As evident from **Figure 5**, the performances demonstrated a relatively high correlation between tasks. Given the substantial impact of layer selection for feature extraction in the evaluated tasks, opting for the feature layer based on pathological finding prediction would serve as a powerful optimization strategy for representation learning models of toxicopathological images, providing deeper insights into the effects of compounds *in vivo*.

**Figure 5.**
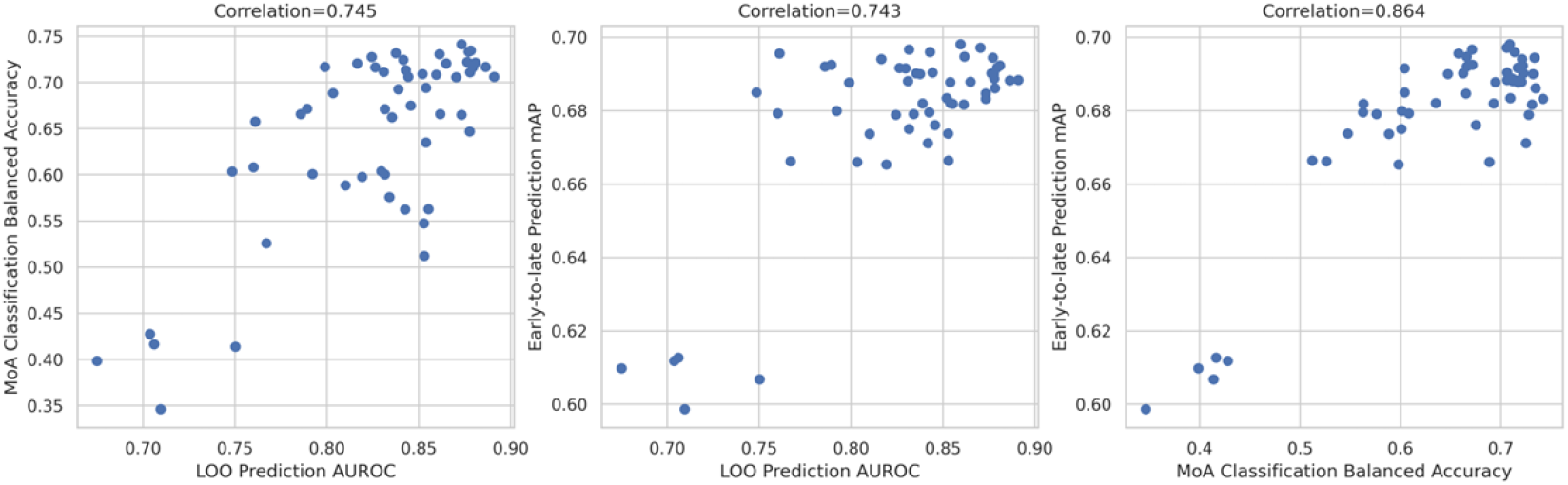
Evaluation of the relationship between the accuracy of estimating pathological findings and downstream tasks. The relationship between the accuracy of estimating pathological findings and downstream tasks is examined. Representative metrics, including AUROC of LOO prediction, Balanced accuracy of MoA classification, and mAP of early-to-late prediction, were selected and visualized as scatter plots. Pearson correlation coefficient is indicated on the top of each panel.

## 5 Conclusion

The contribution of this study is three-fold:

1. We observed that the latent representation of toxicopathological images, extracted by a CNN model, outperforms the direct use of pathological findings in predicting compounds’ properties *in vivo*.
2. The selection of the layer for feature extraction plays a crucial role in obtaining effective representations from toxicopathological images for tasks related to compounds’ effects *in vivo* compared with learning pathological findings.
3. Additionally, we found a positive correlation between pathological finding prediction during the construction of representation learning models and downstream tasks related to compound property estimation, which provides valuable information for feature layer selection.

These findings would serve as a foundation for capturing the biological responses of an organism and numerizing compound effects in vivo by utilizing toxicopathological images routinely obtained in the development of low molecular weight compounds, such as drugs. Appropriate numerization of objects promotes data-driven analysis and would expand what we can recognize and handle about compound properties in vivo, as already shown *in vitro* [35].

While the above direction is relatively focused on basic science, the combination of numerizing toxicopathological images and data analysis would contribute to applied science as well, such as enhancing the detection of latent drug toxicity and improving safety assessment efficiency. For example, Early-to-Late prediction has the potential to reduce the burden of pathological analysis in repeated-dose tests, although further experiments, such as assessing the monotonicity of pathological changes for each finding, are necessary. Expanding beyond liver tissues is essential for enhancing safety assessment efficiency, although the liver is highly descriptive for compound effects due to the first-pass effect [14]. Drawing from Zingman’s work, incorporating multiple tissues would strengthen CNN models and contribute to obtaining effective representations of toxicopathological images, marking an important future direction [13]. Additionally, the expansion of pathological findings is crucial in this direction. Although we limited the pathological findings due to computational cost in handling WSI for an exhaustive comparison of layer dependency, including findings whose WSI-level label is rarely observed at the patch-level using techniques such as multiple instance learning would enhance the ability of the representation learning model to capture compound effects *in vivo*.

## 6 Declarations

### Code & Data Availability

Code, models, and data are available at: https://github.com/mizuno-group/ToxRepresentationCNN.

### Author contributions

Shotaro Maedera: Methodology, Software, Investigation, Writing – Original Draft, Visualization. Tadahaya Mizuno: Conceptualization, Resources, Supervision, Project administration, Writing – Original Draft, Writing – Review & Editing, Funding acquisition.

Hiroyuki Kusuhara: Writing – Review & Editing, Funding acquisition.

## Acknowledgements

We appreciate Dr. Hisamitsu Hayashi (The University of Tokyo) for helpful discussion about this study. We thank all those who contributed to the construction of the following data sets employed in the present study such as Open TG-GATEs and DrugBank. This work was supported by AMED under Grant Number JP22mk0101250h and 22ama121051j0001, the JSPS KAKENHI Grant-in-Aid for Scientific Research (C) (grant number 21K06663) from the Japan Society for the Promotion of Science, and Takeda Science Foundation.

## Supplementary Information

### Supplementary Note. 1: Implementation details of weakly supervised learning for pathological finding selection

For each pathological finding, we divided the positive image experimental groups into 4:1 fractions and similarly divided the negative image experimental groups into 4:1 fractions. Additionally, we divided the experimental groups of negative images, where the finding was absent, in a 4:1 ratio. Images assigned to the 4 sets were used for WSL model training, while the remaining ones were used for validation. Due to the low percentage of positive images in the TGGATEs data, even the largest hypertrophy accounted for only 4.7% of all TGGATEs liver images. To handle data imbalance, we performed under-sampling and randomly cropped 512×512-pixel images from patches, augmenting data through horizontal and vertical flipping.

The cross-entropy loss function was employed, and the EfficientNetB4 model was trained for 20 epochs with a batch size of 32. The initial learning rate was set to 5e-4 and updated every epoch using the cosine annealing scheduler with T_max=10 and eta_min=1e-6. We used the noisy student weight as the starting point for training [1]. Cross entropy was evaluated with a validation set at each epoch, and the model of the best epoch was saved as the WSL model. For each pathological finding, we calculated the probability of a patch extracted from a positive image to contain the pathological finding using the WSL model.

### Supplementary Note. 2: Implementation details of Leave one out (LOO) experiment for testing generalization performance

EfficientNetB4 was trained using the TGGATEs training group and TGGATEs validation group to classify all but one of the seven selected findings out of the eight. The model consists of 9 parts: the convolution stem, 7 invert residual blocks, and the convolution head, with the first k parts frozen, where k varies from 0 to 8. For each pathological finding and each k value, we constructed a model, resulting in a total of 72 models. Training was conducted with five different seeds (123, 124, 125, 126, and 127) to examine result reproducibility.

As patches taken from the same WSI are highly redundant, training with all patches in the TGGATEs training group would converge quickly. To address this, we randomly sampled 1/10 of the patches at each epoch and used them as training data, ensuring a fair comparison of the frozen models. The number of patches extracted was kept constant, regardless of the removed pathological finding and the number of frozen parts when training was performed on the same seed.

For training data preparation, 512×512-pixel images were randomly cropped from patches, and data augmentation was applied through horizontal and vertical flipping. The cross-entropy loss function was used, and the EfficientNetB4 model was trained for 5 epochs with a batch size of 16. The initial learning rate was set to 5e-4 and updated every epoch using the cosine annealing scheduler with T_max=5 and eta_min=1e-6. The noisy student weight served as the starting point for training. Cross-entropy was evaluated with a validation set at each epoch, and the model from the best epoch was saved as the LOO model.

### Supplementary Note. 3: Implementation details of ResNet-based models

ResNet50 with anti-aliasing [2] was trained using the TGGATEs training group and TGGATEs validation group to classify all but one of the seven selected findings out of the eight. The model consists of 5 parts: the convolution stem and 4 convolutional blocks, and we froze the first k parts, where k varies from 0 to 4. For each pathological finding and each k value, we constructed a model, resulting in a total of 40 models. The seed used for ResNet50 training was set as 123.

To address the redundancy in patches taken from the same WSI and to prevent quick convergence with all patches in the TGGATEs training group, we randomly sampled 1/10 of the patches at each epoch and used them as training data. 512×512-pixel images were randomly cropped from patches, and data augmentation was applied through horizontal and vertical flipping. The cross-entropy loss function was employed, and the ResNet50 models were trained for 5 epochs with a batch size of 16. The initial learning rate was set to 5e-4 and updated every epoch using the cosine annealing scheduler with T_max=5 and eta_min=1e-6. We used an ImageNet weight as the starting point for training. Cross-entropy was evaluated with a validation set at each epoch, and the model from the best epoch was saved as the LOO model. All results derived from the ResNet-based models are summarized in **Supplementary Data 2**.

---

**Supplementary Figure. 1:**
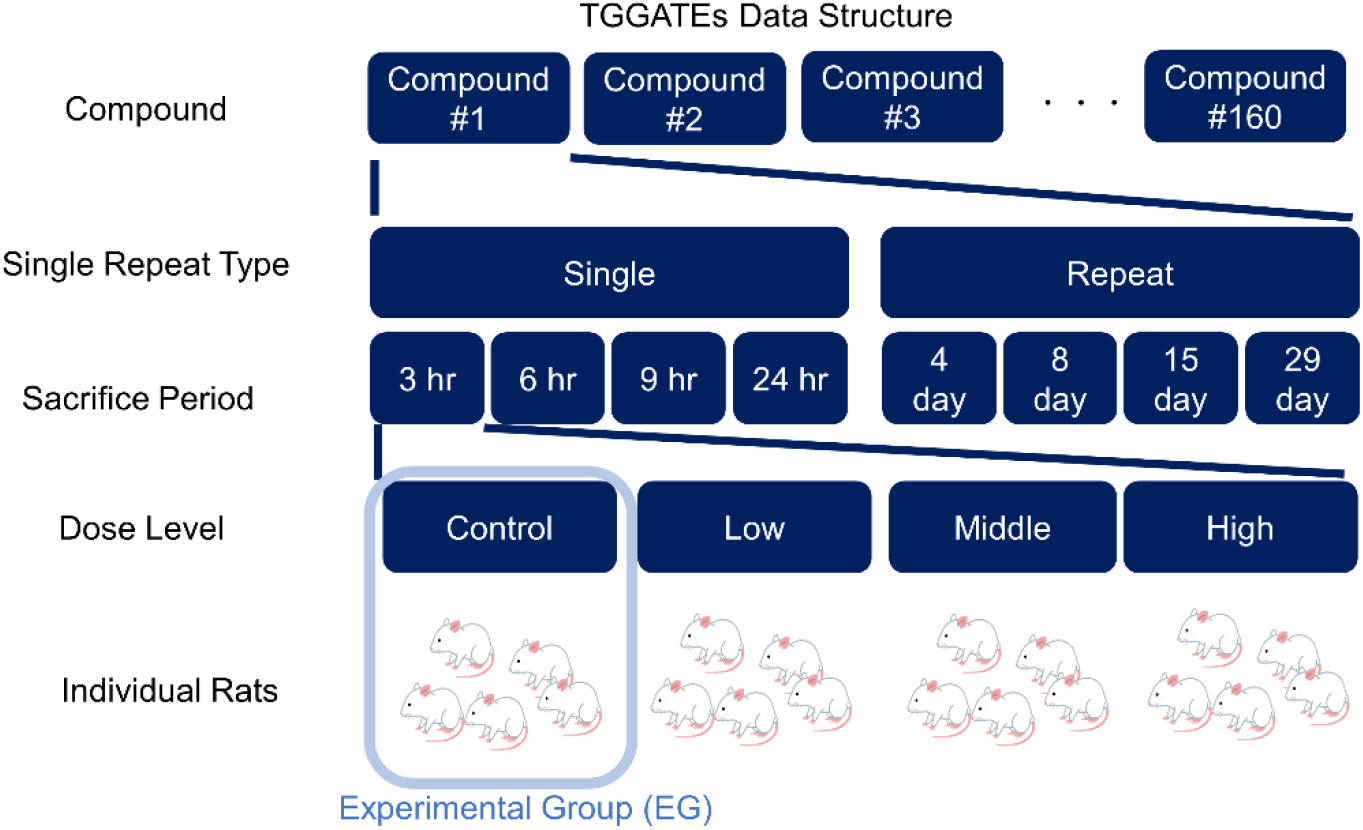
Structure of meta information of Open TG-GATEs data.

**Supplementary Figure. 2:**
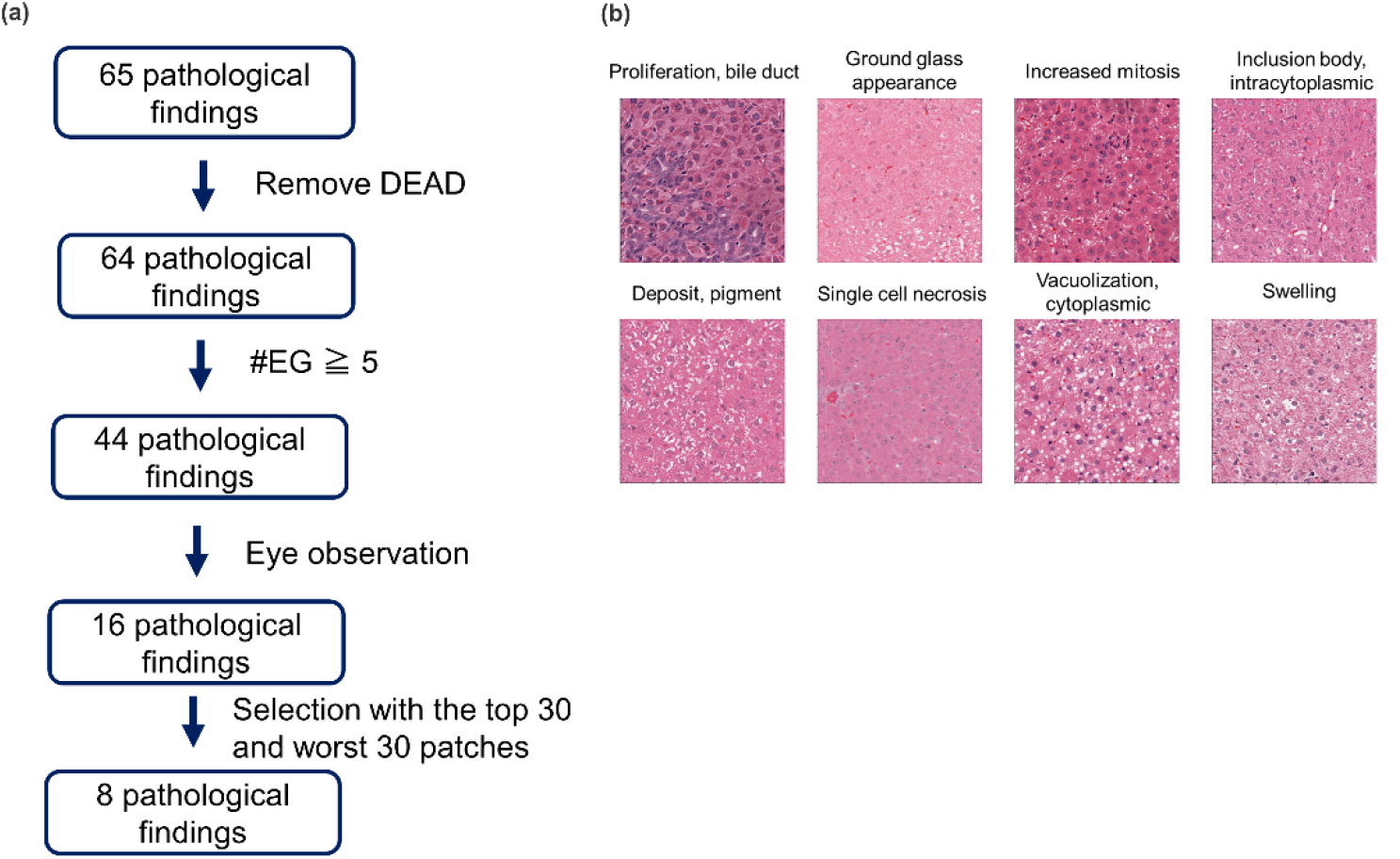
Pathological finding selection utilizing weakly supervised learning. (a) Flowchart depicting the process for narrowing down the pathological findings mainly analyzed in this work. (b) Images showing representative patches of the selected findings.

**Supplementary Figure. 3:**
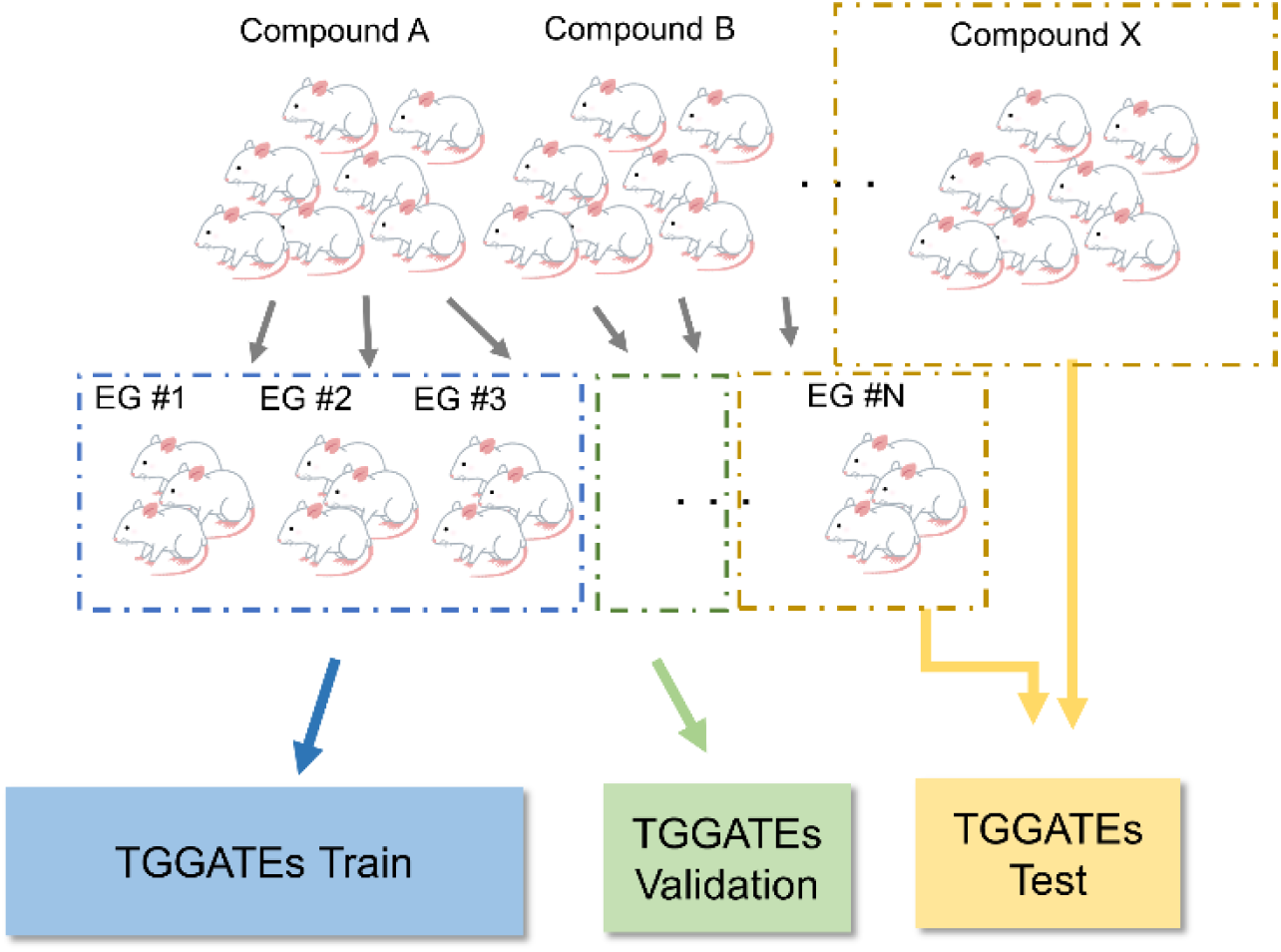
Data-split strategy for Open TG-GATEs images. EG represents Experimental Group, which denotes batch information indicating that the results were obtained in an experiment conducted simultaneously.

**Supplementary Figure 4.**
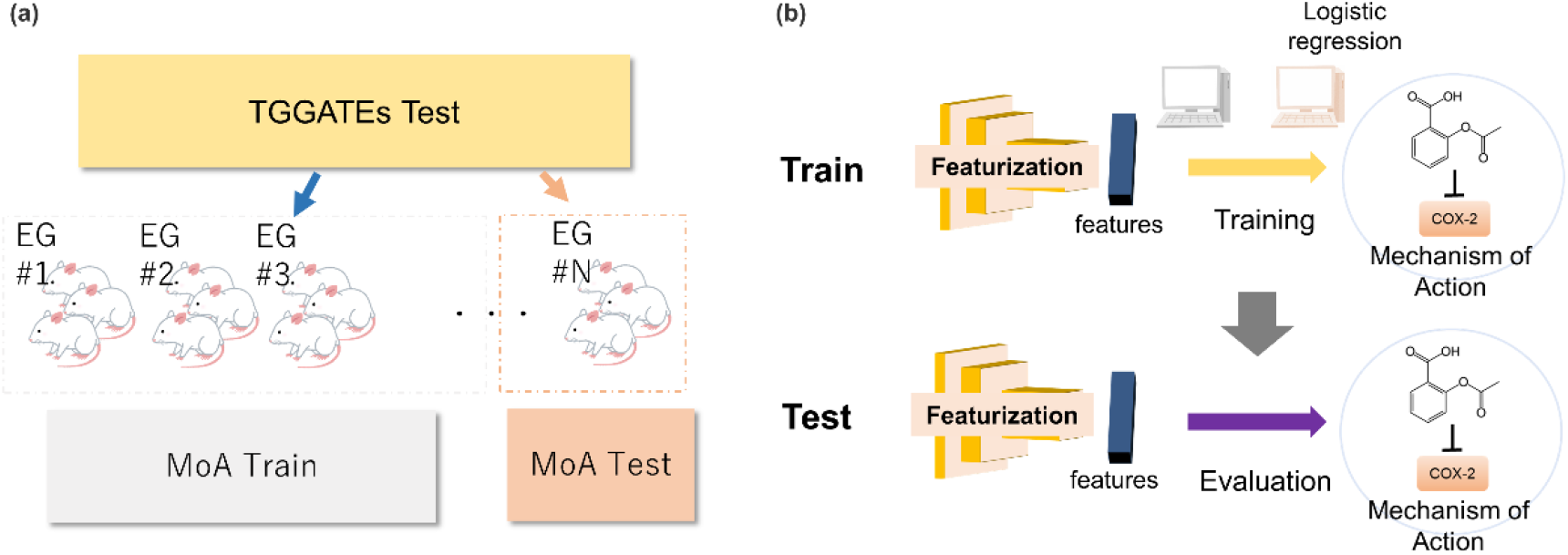
Design of MoA classification. (a) Data-split strategy for MoA classification, with EG representing experimental group. (b) Strategy for modeling and its evaluation.

**Supplementary Figure 5.**
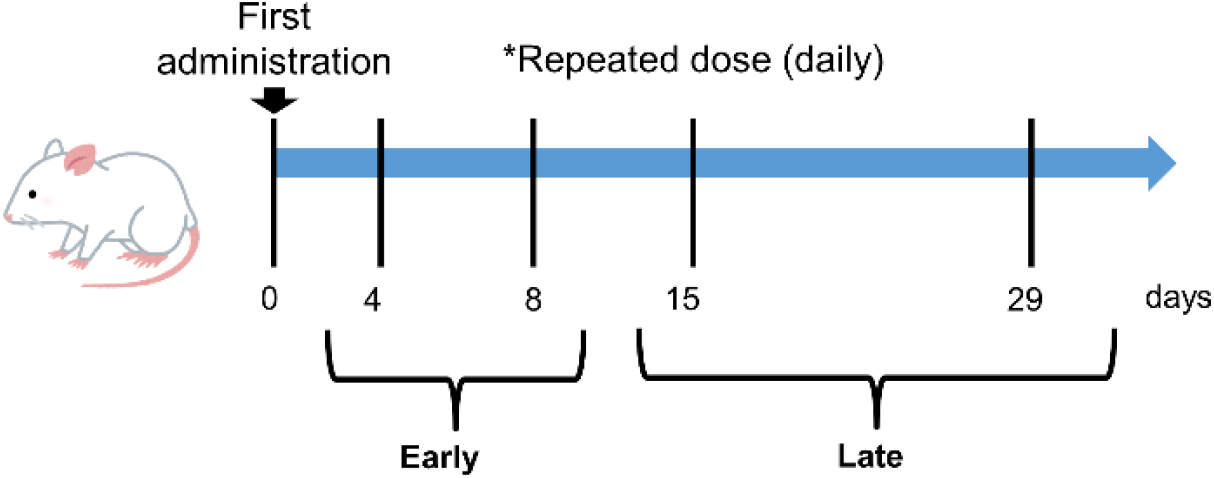
Design of repeated-dose tests in Open TG-GATEs. Daily dosing is conducted until 29 days. The images taken at 4 and 8 days are referred to as the “Early” phase, while the images taken at 15 and 29 days are referred to as the “Late” phase.

**Supplementary Figure 6.**
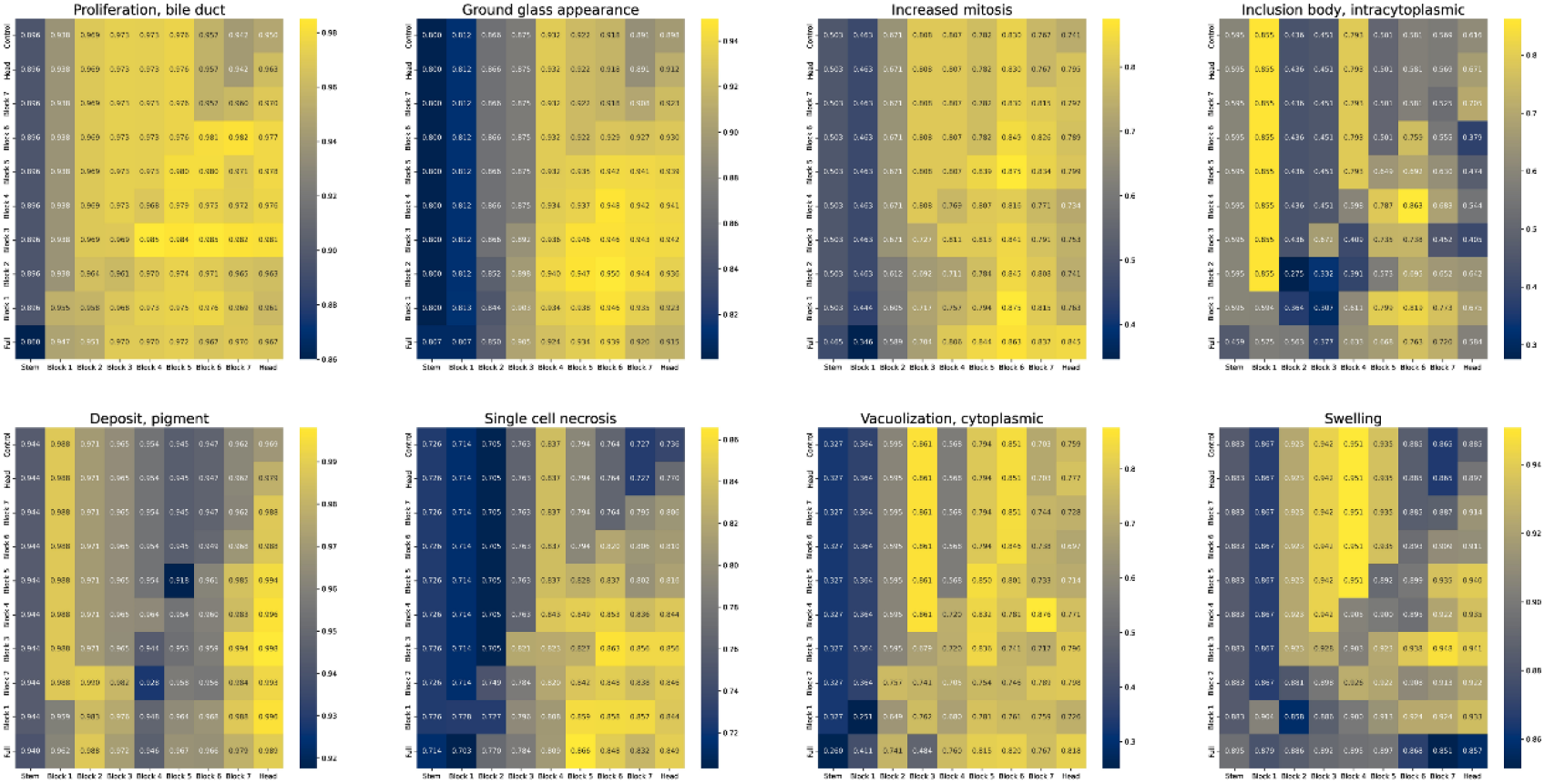
Results of LOO experiments for each finding. AUROC for each finding is presented as a heatmap. The visualization includes the mean values obtained from 5 different seeds.

**Supplementary Figure 7:**
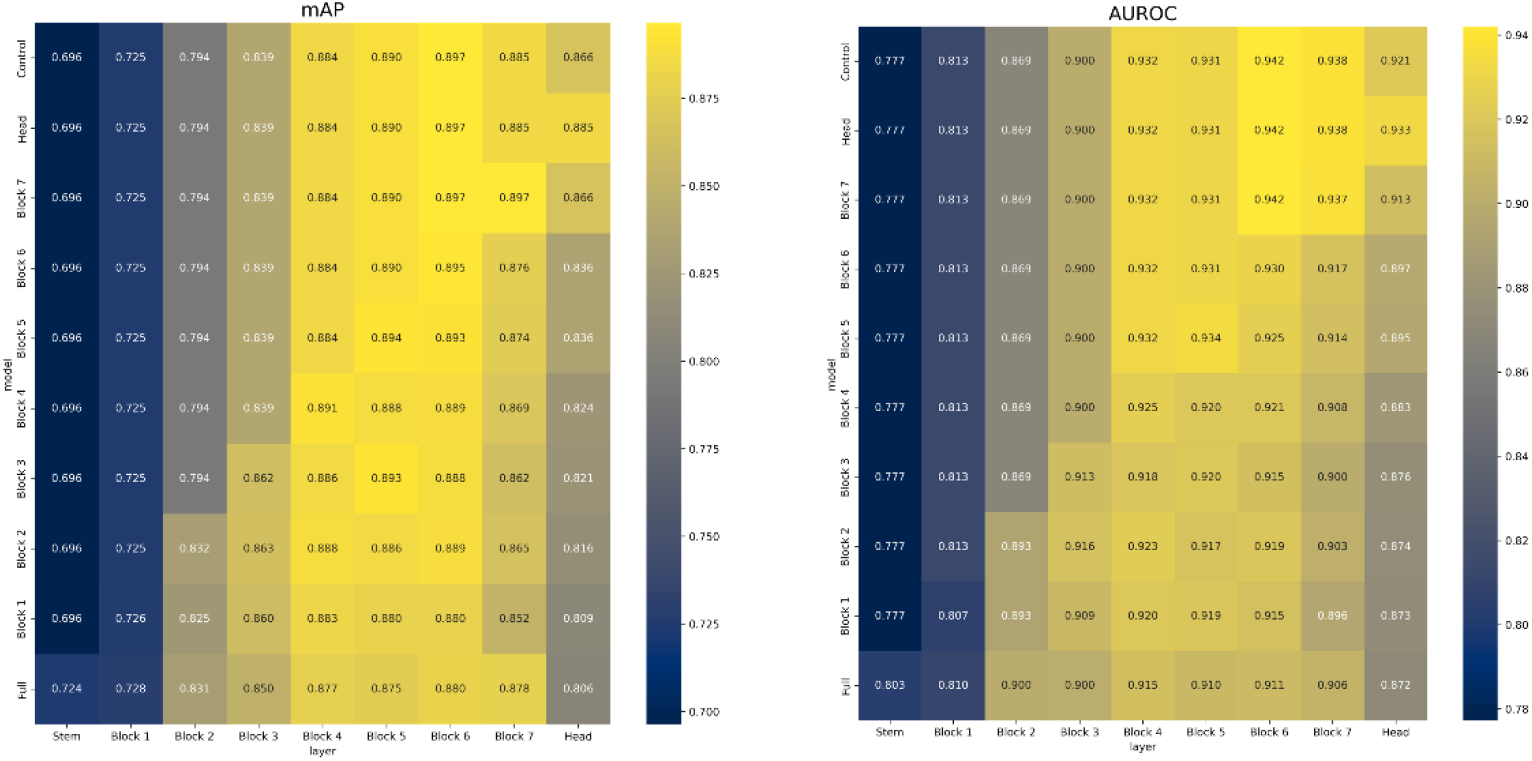
mAP of MoA classification. mAP of each MoA classification is calculated and visualized as a heatmap. The heatmap displays the mean values obtained from 5 different seeds.

**Supplementary Figure 8:**
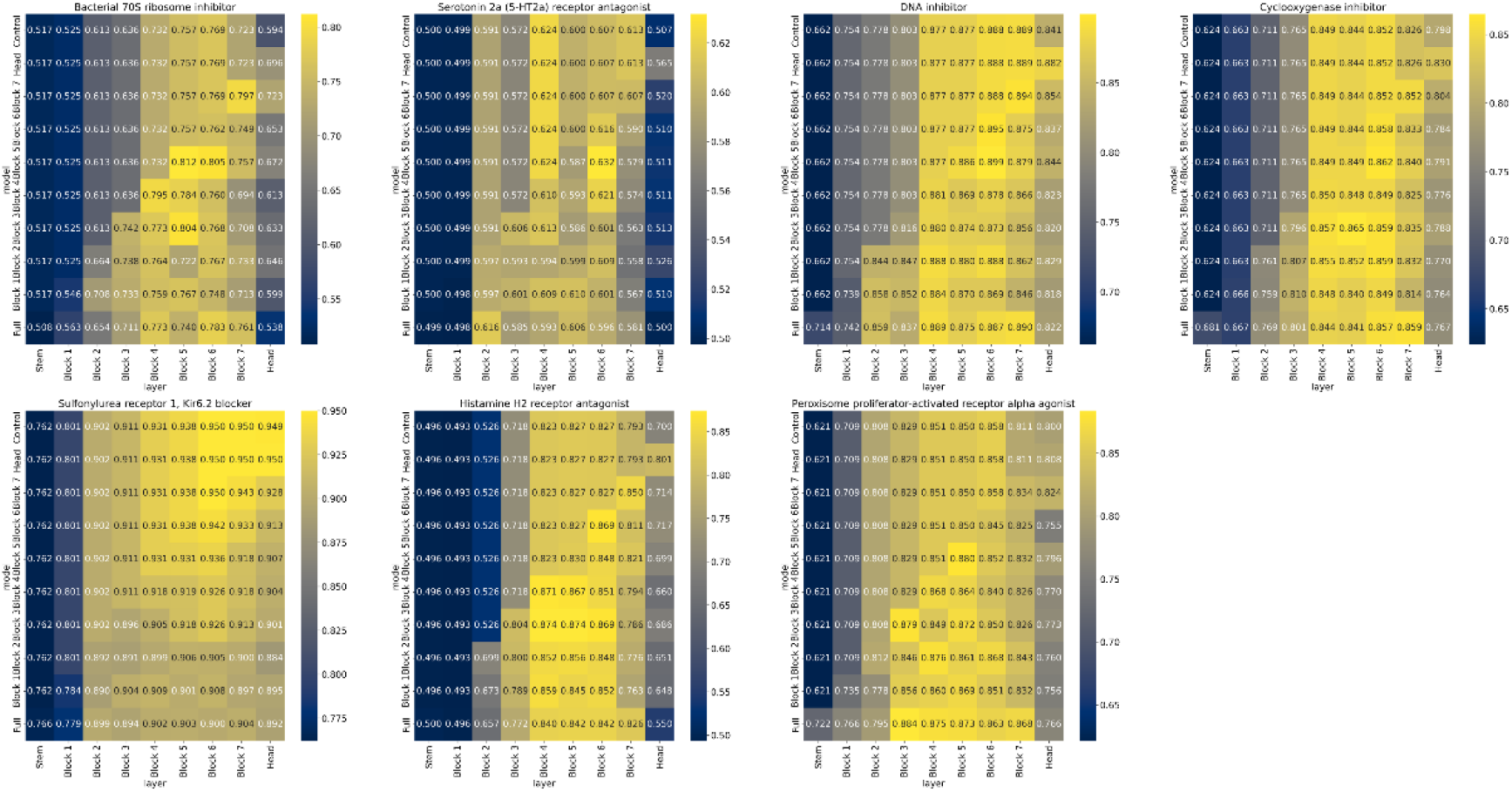
Results of MoA classification for each MoA. AUROC for each MoA is visualized as a heatmap, displaying the mean values obtained from 5 different seeds.

**Supplementary Figure 9:**
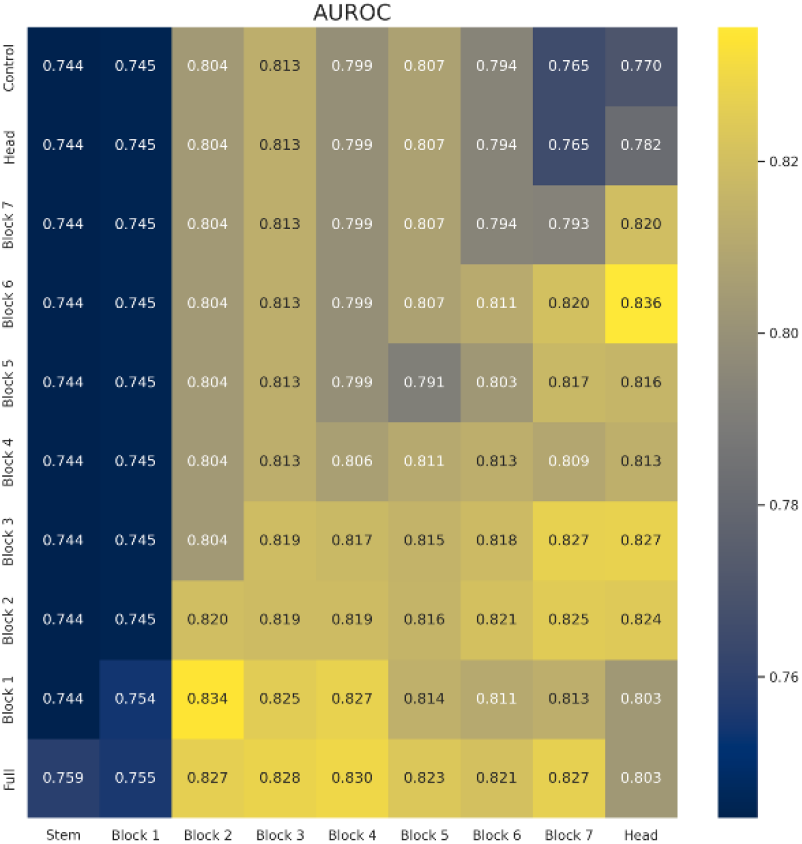
AUROC of Early-to-late prediction of findings. AUROC of each finding prediction is calculated and presented as a heatmap, visualizing the mean values obtained from 5 different seeds.

**Supplementary Table 1:** List of pathological findings in Open TG-GATEs.

| ID | Pathological Finding | #Positive Image | Positive Ratio |
| --- | --- | --- | --- |
| 1 | Alteration, cytoplasmic | 36 | 0.001506213 |
| 2 | Alteration, nuclear | 61 | 0.002552194 |
| 3 | Anisonucleosis | 52 | 0.002175641 |
| 4 | Atrophy | 12 | 0.000502071 |
| 5 | Cellular foci | 31 | 0.001297017 |
| 6 | Cellular infiltration | 482 | 0.02016652 |
| 7 | Cellular infiltration, mononuclear cell | 72 | 0.003012426 |
| 8 | Change, acidophilic | 110 | 0.004602318 |
| 9 | Change, basophilic | 70 | 0.002928748 |
| 10 | Change, eosinophilic | 398 | 0.016652023 |
| 11 | Degeneration | 24 | 0.001004142 |
| 12 | Degeneration, acidophilic, eosinophilic | 19 | 0.000794946 |
| 13 | Degeneration, fatty | 88 | 0.003681854 |
| 14 | Degeneration, granular, eosinophilic | 237 | 0.009915903 |
| 15 | Degeneration, hydropic | 56 | 0.002342998 |
| 16 | Deposit, glycogen | 67 | 0.00280323 |
| 17 | Deposit, hemosiderin | 17 | 0.000711267 |
| 18 | Deposit, pigment | 40 | 0.00167357 |
| 19 | Dilatation | 48 | 0.002008284 |
| 20 | Edema | 93 | 0.003891051 |
| 21 | Fibrosis | 69 | 0.002886908 |
| 22 | Granuloma | 14 | 0.00058575 |
| 23 | Ground glass appearance | 210 | 0.008786243 |
| 24 | Hematopoiesis, extramedullary | 69 | 0.002886908 |
| 25 | Hemorrhage | 54 | 0.00225932 |
| 26 | Hypertrophy | 1134 | 0.047445714 |
| 27 | Inclusion body, intracytoplasmic | 53 | 0.00221748 |
| 28 | Increased mitosis | 428 | 0.017907201 |
| 29 | Inflammation | 7 | 0.000292875 |
| 30 | Lesion,NOS | 44 | 0.001840927 |
| 31 | Microgranuloma | 447 | 0.018702146 |
| 32 | Mineralization | 7 | 0.000292875 |
| 33 | Necrosis | 614 | 0.025689302 |
| 34 | Nodule, hepatodiaphragmatic | 13 | 0.00054391 |
| 35 | Proliferation | 6 | 0.000251036 |
| 36 | Proliferation, Kupffer cell | 50 | 0.002091963 |
| 37 | Proliferation, bile duct | 161 | 0.00673612 |
| 38 | Proliferation, oval cell | 66 | 0.002761391 |
| 39 | Pyknosis | 33 | 0.001380695 |
| 40 | Scar | 6 | 0.000251036 |
| 41 | Single cell necrosis | 445 | 0.018618468 |
| 42 | Swelling | 246 | 0.010292456 |
| 43 | Vacuolization, cytoplasmic | 317 | 0.013263043 |
| 44 | Vacuolization, nuclear | 15 | 0.000627589 |

**Supplementary Table 2:** List of mechanism of action in Open TG-GATEs.

| ID | Mechanism of Action | Name of Compounds | Focused or not |
| --- | --- | --- | --- |
| 1 | Cyclooxygenase inhibitor | naproxen; sulindac; sulfasalazine; diclofenac; acetaminophen; phenylbutazone; mefenamic acid; indomethacin; aspirin | Yes |
| 2 | DNA inhibitor | nitrofurantoin; carboplatin; azathioprine; nitrofurazone; lomustine; cyclophosphamide | Yes |
| 3 | Bacterial 70S ribosome inhibitor | erythromycin ethylsuccinate; chloramphenicol; tetracycline | Yes |
| 4 | Histamine H2 receptor antagonist | cimetidine; famotidine; ranitidine | Yes |
| 5 | Peroxisome proliferator-activated receptor alpha agonist | fenofibrate; gemfibrozil; clofibrate | Yes |
| 6 | Serotonin 2a (5-HT2a) receptor antagonist | chlorpromazine; thioridazine; haloperidol | Yes |
| 7 | Sulfonylurea receptor 1, Kir6.2 blocker | glibenclamide; tolbutamide; chlorpropamide | Yes |
| 8 | Thyroid peroxidase inhibitor | methimazole; propylthiouracil | No |
| 9 | Calmodulin inhibitor | benziodarone | No |
| 10 | Histamine H1 receptor antagonist | hydroxyzine; promethazine | No |
| 11 | Xanthine dehydrogenase inhibitor | allopurinol; bendazac | No |
| 12 | Aldehyde dehydrogenase inhibitor | disulfiram | No |
| 13 | Cytochrome P450 51 inhibitor | ketoconazole | No |
| 14 | D2-like dopamine receptor antagonist | chlorpromazine; thioridazine | No |
| 15 | Androgen Receptor antagonist | flutamide | No |
| 16 | Glucocorticoid receptor agonist | dexamethasone | No |
| 17 | Squalene monooxygenase inhibitor | terbinafine | No |
| 18 | Enoyl-[acyl-carrier-protein] reductase inhibitor | isoniazid; ethionamide | No |
| 19 | Sodium channel alpha subunit blocker | phenytoin; carbamazepine | No |
| 20 | Adrenergic receptor alpha antagonist | moxisylyte | No |

**Supplementary Table 2** continued.
|  |  |  |  |
| --- | --- | --- | --- |
| 21 | Voltage-gated L-type calcium channel blocker | nifedipine | No |
| 22 | Arachidonate 5-lipoxygenase inhibitor | sulfasalazine | No |
| 23 | Tubulin inhibitor | griseofulvin; colchicine | No |
| 24 | Amidophosphoribosyltransferase inhibitor | azathioprine | No |
| 25 | 80S Ribosome inhibitor | cycloheximide | No |
| 26 | Potassium-transporting ATPase inhibitor | omeprazole | No |
| 27 | Anandamide amidohydrolase inhibitor | acetaminophen | No |
| 28 | Vanilloid receptor opener | acetaminophen | No |
| 29 | Androgen Receptor agonist | danazol; methyltestosterone | No |
| 30 | Progesterone receptor agonist | danazol | No |
| 31 | Succinate semialdehyde dehydrogenase inhibitor | valproic acid | No |
| 32 | Copper chelating agent | penicillamine | No |
| 33 | Voltage-gated T-type calcium channel blocker | trimethadione | No |
| 34 | Serotonin transporter inhibitor | fluoxetine hydrochloride | No |
| 35 | HMG-CoA reductase inhibitor | simvastatin | No |
| 36 | Cyclooxygenase-2 inhibitor | meloxicam | No |
| 37 | Carbonic anhydrase I inhibitor | acetazolamide | No |
| 38 | Carbonic anhydrase II inhibitor | acetazolamide | No |
| 39 | Carbonic anhydrase IV inhibitor | acetazolamide | No |
| 40 | Carbonic anhydrase XII inhibitor | acetazolamide | No |

**Supplementary Table 2** continued.
|  |  |  |  |
| --- | --- | --- | --- |
| 41 | Estrogen receptor alpha agonist | ethinylestradiol | No |
| 42 | Bacterial DNA gyrase inhibitor | ciprofloxacin | No |
| 43 | Norepinephrine transporter inhibitor | adapin | No |
| 44 | Dopamine D2 receptor antagonist | fluphenazine; sulpiride | No |
| 45 | Adenosine receptor antagonist | theophylline; caffeine | No |
| 46 | Phosphodiesterase 3 inhibitor | theophylline | No |
| 47 | Phosphodiesterase 4 inhibitor | theophylline | No |
| 48 | Amiloride-sensitive sodium channel, ENaC blocker | triamterene | No |
| 49 | DNA topoisomerase II inhibitor | etoposide | No |
| 50 | Vasopressin receptor agonist | desmopressin acetate | No |
| 51 | 26S proteasome inhibitor | bortezomib | No |
| 52 | HERG blocker | amiodarone | No |
| 53 | Angiotensin-converting enzyme inhibitor | captopril | No |
| 54 | Type I iodothyronine deiodinase (Type-I 5'-deiodinase) (DIOI) (Type 1 DI) (5DI) inhibitor | propylthiouracil | No |
| 55 | GABA-A receptor | diazepam; triazolam | No |
| 56 | anion channel positive allosteric modulator | diazepam; triazolam | No |
| 57 | Solute carrier family 22 member 12 inhibitor | benzbromarone | No |
| 58 | Serotonin 2c (5-HT2c) receptor antagonist | thioridazine | No |
| 59 | D2-like dopamine receptor inverse agonist | haloperidol | No |
| 60 | Epidermal growth factor receptor erbB1 inhibitor | gefitinib | No |
| 61 | Maltase-glucoamylase inhibitor | acarbose | No |
| 62 | Pancreatic alpha-amylase inhibitor | acarbose | No |
| 63 | Peroxisome proliferator-activated receptor gamma agonist | rosiglitazone maleate | No |
| 64 | Bacterial DNA-directed RNA polymerase inhibitor | rifampicin | No |
| 65 | Bacterial penicillin-binding protein inhibitor | cephalothin | No |
| 66 | Monoamine oxidase inhibitor | iproniazid | No |
| 67 | Reducing agent | tiopronin | No |

